# Host insect microRNAs are translocated to an obligate endosymbiont

**DOI:** 10.64898/2026.07.17.739287

**Authors:** Yaping Ding, Hongyan Sun, Georg Jander, Alex C.C. Wilson, Honglin Feng

**Author notes:** **Author for Correspondence:** Honglin Feng, Department of Entomology, Louisiana State University AgCenter, Baton Rouge, LA 70803, USA.

## Abstract

Many insects rely on intimate interactions with bacterial symbionts housed in specialized cells called bacteriocytes. While host and symbiont gene expression in bacteriocytes appears highly integrated, the underlying regulatory mechanisms remain largely unknown. MicroRNAs regulate gene expression and are emerging as mediators of cross-kingdom communication. Here, in the pea aphid, *Acyrthosiphon pisum*, and the green peach aphid, *Myzus persicae*, we demonstrate cross-kingdom translocation of aphid miRNAs into their obligate endosymbiont, *Buchnera aphidicola*. miRNA fluorescence *in situ* hybridization in aphid embryos provides direct evidence that four out of five candidate miRNAs (miR-1, miR-10, miR-29, and miR-927) localize inside *Buchnera* cells. To investigate how these miRNAs may regulate *Buchnera* gene expression, we further demonstrated the cross-kingdom translocation of aphid argonuate 1 protein (Ago1), into *Buchnera* using immunolocalization. Given the translocation of both Ago1 and miRNAs, we applied a eukaryotic miRNA target prediction framework and found that all cross-kingdom-translocated miRNAs are predicted to target *Buchnera* genes involved in symbiotic functions. Although the precise *in vivo* functions of translocated miRNAs remain challenging to determine, our findings suggest a previously unrecognized layer of regulation between insect host and its obligate endosymbiont, offering new insight into the molecular dialogue that supports insect-microbe interactions, highlighting potential targets for miRNA-based pest management.

**Significance statement:** Many insects have evolved bacteriocyte cells to house essential bacterial endosymbionts. However, how bacteriocytes are specified and how gene expression is coordinated between host and symbiont remain largely unknown. Here, we identify that aphid microRNAs, along with an argonaute protein, are translocated into the obligate bacterial endosymbiont *Buchnera*, revealing a previously unrecognized regulatory layer mediating host and obligate endosymbiont interactions. This discovery provides evidence of miRNA cross-kingdom movement into an obligate endosymbiont, filling a key knowledge gap in how eukaryotic host small RNAs can influence microbial partners. Given the critical role of *Buchnera* in aphid survival and reproduction, these findings not only advance fundamental knowledge of insect-microbe interactions but also point to new molecular targets for innovative pest management strategies.

## Introduction

Many insects, including Hemiptera and some Diptera, form symbiotic relationships with bacteria, housing them in bacteriocytes (1). These specialized host cells are typically highly polyploid and exhibit distinct host transcriptional profiles (2, 3) that are complemented by gene expression from the much smaller symbiont genomes (4–6), to facilitate and maintain the symbiotic relationship. Although both host and symbiont genes are known to be dynamically regulated during bacteriocyte development, differentiation, and maintenance of bacterial symbionts (7–9), the underlying molecular regulatory mechanisms remain poorly understood.

MicroRNAs (miRNAs) are small non-coding RNAs (typically 21-25 nucleotides long) that regulate gene expression at the post-transcriptional level through sequence complementarity (10). They are synthesized in the eukaryotic nucleus from endogenous genes by RNA polymerase II as primary miRNA (pri-miRNA), which are processed by the Drosha-Pasha complex into hairpin-shaped precursor miRNAs (pre-miRNAs). Pre-miRNAs are exported by exportin-5 into the cytoplasm, where the endonuclease Dicer-1 trims the hairpin loop, generating a miRNA duplex. One strand of this duplex becomes the mature miRNA and is loaded onto the RNA-Induced Silencing Complex (RISC) to guide the RISC Central protein, Argonaute, targeting mRNA for gene expression regulation (10).

In insects, miRNAs influence processes such as growth and development (11), immunity (12), insecticide resistance (13), interactions with host plants (14), and symbiosis with bacterial endosymbionts (15, 16). Beyond regulating gene expression within endogenous genomes, miRNAs have also emerged as mediators of cross-kingdom communication. Such cross-kingdom miRNA activity has been documented in diverse systems, including host-pathogen interactions (*e.g*., plant and fungal pathogens) (17), host-facultative symbiont interactions (*e.g*., plant and mycorrhizal fungi) (18), and even interactions between host cells and their organelles, such as mitochondria (19). Despite these advances, it remains unknown whether miRNAs can mediate cross-kingdom regulation in the intimate and ancient symbioses that are fundamental to the biology and evolution of many insects.

Our previous research with the pea aphid, *Acyrthosiphon pisum,* and the green peach aphid, *Myzus persicae*, revealed that some aphid-derived miRNAs are highly enriched in bacteriocytes that harbor the obligate endosymbiont *Buchnera aphidicola* (20). Among these bacteriocyte miRNAs, miR-92a and miR-3024, have been suggested to play roles in aphid bacterial symbiosis. MiR-92a influences the expression of a bacteriocyte-specific gene (21) that may be involved in integration with *Buchnera* during embryonic development (22), while miR-3024 is unique to aphids, regulating nutrient homeostasis between aphids and their secondary endosymbionts (15). These bacteriocyte miRNAs have been shown to regulate host aphid genes involved in the symbiotic interactions (15, 21). However, whether they also regulate symbiont genes via a cross-kingdom mechanism (23) requires investigation. To explore this possibility, we determined whether aphid miRNAs are translocated into *Buchnera*.

## Results

### Aphid miRNAs localize inside Buchnera cells

Based on our earlier work, which identified 14 miRNAs significantly enriched and/or upregulated in the aphid bacteriome relative to the gut (20), and a new annotation of aphid miRNAs in publicly available *Buchnera* small RNAseq datasets (9) (Table S1), we selected five aphid miRNAs (miR-1, miR-10, miR-29, miR-306, and miR-927) and evaluated their potential for cross-kingdom translocation from the aphid host into *Buchnera* using miRNA fluorescent *in situ* hybridization (FISH) (**Fig. 1 and Fig. S1-S6**). We conducted the miRNA FISH experiments in two aphid species, the pea aphid, *Acyrthosiphon pisum,* and the green peach aphid, *Myzus persicae*. Experiments for each miRNA in each species were independently repeated 2-3 times. To establish both the integrity of the miRNA FISH protocol and the specificity of the miRNA probes, all experiments included the following controls: (i) a positive control targeting the ubiquitously expressed small nuclear RNA U6 (24); (ii) a pre-adsorbed control, in which miRNA probes were incubated with a fourfold excess of their miRNA mimic to competitively block probe binding to endogenous miRNAs; and (iii) a no-probe control to exclude nonspecific and background fluorescence. Data collection targeted embryos following cellularization when the *Buchnera* population has been compartmentalized into bacteriocytes (developmental stages 14-15) (7, 25). For each miRNA, a minimum of 10 embryos were imaged per experiment. The number of embryos displaying the reported localization pattern, together with the total number of embryos examined, is reported in the bottom-right corner of all embryo panels (**Fig. 1**). To facilitate the characterization of miRNA localization patterns, we performed line-scan fluorescence intensity analyses across individual *Buchnera* cells. For each miRNA, the individual *Buchnera* average intensity profiles were generated from 50 randomly selected *Buchnera* across all embryos and experiments (**Fig. 1**, column ^v^ - *Intensity*). Line-scan intensities were also analyzed across multiple *Buchnera* cell spaces to facilitate distinction of bacteriocyte cytoplasmic localization (**Fig. S6**). As expected, miRNA U6 was detected in all aphid cells across all embryos examined (**Fig. 1A-A^v^**). Within bacteriocytes, U6 signal was localized to both the cytoplasm and the nucleolus. Notably, although its function remains unclear, we cannot exclude the possibility that U6 is translocated into *Buchnera*, as U6 signals were detected within some *Buchnera* cells (**Fig. 1A^ii^ and Fig. S6**).

**Figure 1.**
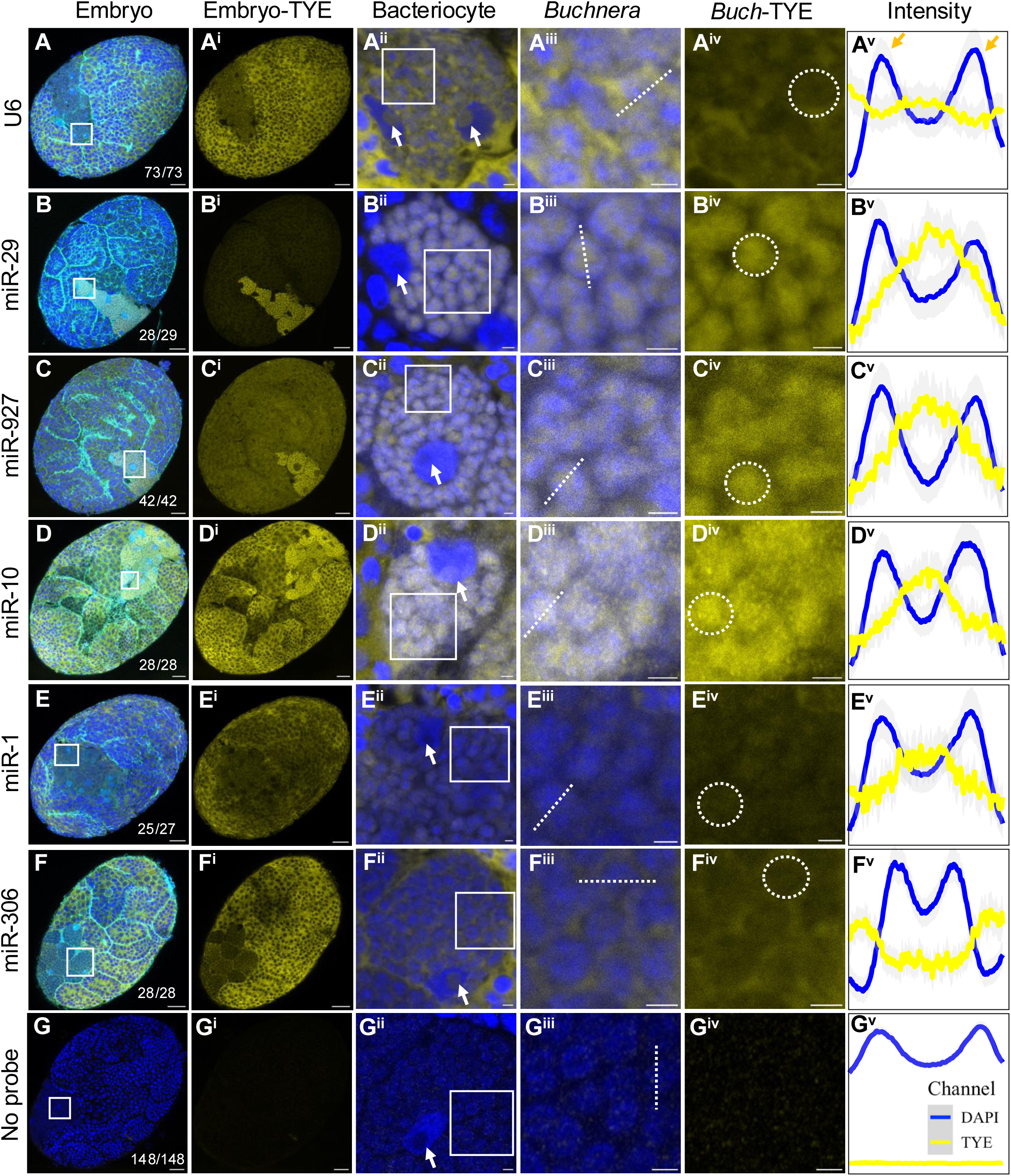
miRNA localization in *A. pisum* embryo, bacteriocyte, and *Buchnera* using fluorescent miRNA *in situ* hybridization. All embryos are in stages 14-15 with bacteriocyte cells clearly formed. Blue indicates DAPI staining of DNA; Cyan indicates phalloidin staining of F-actin, only shown in the first column (Embryo); Yellow indicates miRNA localization visualized using the TYE665 fluorophore. Squares mark regions shown at higher magnification in the adjacent columns. Dashed circles outline individual *Buchnera* cells. The intensity panel (“**v**”) show line-scan average intensity measurements of 50 randomly selected *Buchnera* cells across different embryos and across all experiments. Blue traces indicate DAPI intensities with the two major peaks, and yellow traces indicate TYE665 miRNA signal intensity. Shaded regions along the traces indicate the 95% confidence interval. Orange arrows indicates the cytoplasmic miRNA signals. Each miRNA FISH experiment with all controls was repeated at least twice across *A. pisum* and *M. persicae*. The number of embryos showing the observed patterns out of the total are summarized in the bottom right corner (**A–G**). All images were taken under the same confocal settings. Scale bar represents 20 µm in panels with embryos, and 2 µm in panels with bacteriocytes or *Buchnera* cells. **(A–A^v^)** U6, a ubiquitous small nuclear RNA, is enriched in the nucleolus of all aphid cells (A-A^iii^), and its signal in bacteriocyte is localized to the bacteriocyte nucleus (A^ii^) and cytoplasm (A^iv-^A^v^). **(B–B^v^)** miR-29 and **(C–C^v^)** miR-927 are significantly enriched in bacteriocytes (B^i^ & C^i^). Higher magnification (B^ii^-B^iv^ & C^ii^-C^iv^) reveals that these miRNAs localize within *Buchnera* cells (dashed circles), with clear signal detected between the *Buchnera* boundaries (B^v^ and C^v^). **(D–D^v^)** miR-10 is enriched in bacteriocytes (D-D^iii^) and localized within *Buchnera* cells (D^v^), displaying a more diffuse pattern (D^iv^). (**E–E^v^**) miR-1 shows low expression in bacteriocytes (E-E^iv^) but detectable signal is present within *Buchnera* cells (E^iv^-E^v^). **(F– F^v^)** miR-306 is enriched in other tissues compared to bacteriocytes in the developing embryos (F-F**^ii^**). Although present in bacteriocytes (F^ii^), its signal is localized to the cytoplasm and not within *Buchnera* (F^iii^-F^v^). (**G-G^v^**) no specific signals were detected in no-probe controls across all experiments. Additionally, no specific signals were detected in pre-adsorbed controls that are specific to each miRNAs (**Figure S1-S5**).

Aphid miRNAs revealed two distinct patterns of localization in embryonic bacteriomes. Four of the five tested miRNAs (miR-29, miR-927, miR-10, and miR-1) were found to be localized inside *Buchnera* cells (**Fig. 1** rows B-E). By contrast, the fifth miRNA, miR-306, was found to localize to the cytoplasm of bacteriocytes. Notably, two of the four *Buchnera*-localized miRNAs (miR-29 & miR-927) were enriched in bacteriocytes relative to other embryonic tissues (**Fig. 1 B^i^ & C^i^**). Within bacteriocytes, miR-29 exhibited strong and well-defined localization within *Buchnera* cells (**Fig. 1B^ii^^-iv^**) with line-scan intensity analysis revealing a strong miR-29 signal peak between the DAPI-defined *Buchnera* cell boundary (**Fig. 1B^v^, Fig. S6**). While the miR-927 signal was also strongly enriched in bacteriocytes relative to other embryonic tissues (**Fig. 1C-C^i^**), its signal in non-bacteriocyte tissues (**Fig. 1C^i^**) was stronger than that observed for miR-29 (**Fig. 1B^i^**). Within bacteriocytes, miR-927 exhibited strong and well-defined signals within *Buchnera* cells (**Fig. 1C^ii^-C^v^**), with the line-scan intensity profile confirming its enrichment within DAPI-defined *Buchnera* cell boundaries (**Fig. 1C^v^, Fig. S6**). In contrast to miR-29 and miR-927, miR-10 and miR-1 were strongly expressed across almost all embryonic cells (**Fig. 1 D^i^ & E^i^**). While miR-10 signal was detected within *Buchnera* cells (**Fig. 1 D^ii-v^)**, its localization pattern appeared more punctate than that observed for miR-29 and miR-927 (**Fig. 1D^iii^^-iv^**). By contrast, miR-1 displayed a high expression in non-symbiotic tissues compared to bacteriocytes (**Fig. 1E-E^i^**). Despite its relatively low abundance in bacteriocytes (**Fig. 1E^ii^-E^iv^**), line-scan intensity analyses detected miR-1 signals within *Buchnera* cells (**Fig. 1E^v^, Fig. S6**). Like miR-1, miR-306 was enriched in non-symbiotic tissues relative to bacteriocytes (**Fig. 1F-F^i^**). However, unlike the four miRNAs described above, miR-306 fluorescence signal within the bacteriocytes was localized to the cytoplasmic region surrounding *Buchnera* rather than within *Buchnera* cells (**Fig. 1F^ii^-F^v^**). This distinct localization pattern reinforces the specificity of the cross-kingdom localization observed for miR-29, miR-927, miR-10, and miR-1. Replication of these miRNA FISH experiments in stage 14-15 *M. persicae* embryos revealed localization patterns identical to those described above for *A. pisum* (**Fig. S1-S5**).

With respect to the controls in all experiments, no specific signal was detected in either the pre-absorbed or no-probe controls for all miRNAs (**Fig. 1G-G^v^**), thereby confirming probe specificity. The pre-absorbed controls were performed for each miRNA independently and are not shown here for *A. pisum* due to figure space limitations, but the results were consistent with those pre-absorbed controls observed in *M. periscae* and presented in **Figs. S1-S5**.

To further explore the taxonomic breadth of cross-kingdom translocation of aphid RNA to *Buchnera*, we mined existing small RNAseq datasets representative of five aphid species from three genera (*Acyrthosiphon*, *Uroleucon*, and *Schizaphis*) within the family of Aphididae (9). We found that miR-1 and miR29 are not only consistently detected across all *Buchnera* small RNAseq datasets, but also significantly enriched in isolated *Buchnera* compared with bacteriocyte (**Table S1**). Finding miR-1 and miR-29 in these *Buchnera* small RNA datasets suggest that miRNA translocation into *Buchnera* is conserved and ancient.

### Aphid argonaute 1 protein localized to Buchnera cells

miRNAs regulate gene expression through the miRNA pathway involving the Argonaute (Ago) protein, the core functional component of the RNA-induced silencing complex (RISC) (26). Although multiple Ago protein families exist, in insects, the Ago1-RISC complex has been shown to primarily mediate miRNA-guided post-transcriptional gene regulation (27, 28). In aphids, the Ago1 family comprises two paralogs, Ago1a and Ago1b, where Ago1b is a duplicated variant that has undergone positive selection (29, 30). In *A. pisum*, Ago proteins exhibit morph- and stage-specific expression (30, 31). Particularly, for Ago1, both Ago1a and Ago1b are highly expressed in parthenogenetic sexuparae and sexual oviparous females and males, whereas Ago1a is more highly expressed than Ago1b in parthenogenetic virginoparous females (30). Prior to this study, the tissue-specific expression of Ago1a and 1b has not been characterized.

Given the central role of Ago1 in miRNA-mediated gene silencing, we sought to determine its localization in aphid bacteriocytes by generating an antibody. We selected *A. pisum* Ago1a for antibody production—aphid Ago1 shows high sequence and structural similarity among isoforms and across *A. pisum* and *M. persicae* (**Fig. S7A**),. For antibody generation, the full-length Ago1a protein was expressed in *E. coli* BL21 Star^TM^(DE3) using the pET30a expression vector; and the purified recombinant protein was used to immunize mice through three consecutive injections. Antibody purification was validated using ELISA, and its specificity was confirmed by Western blot using aphid protein extracts, with the recombinant protein serving as a positive control (**Fig. S7B**). When total protein was extracted from more than 300 mg of whole aphids, Western blot analysis using Ago1a antibody detected a prominent band corresponding to the predicted molecular weight of Ago1a, and a weaker low-abundant band corresponding to Ago1b (**Fig. S7B**). Mass spectrometry analysis of these bands identified peptides derived from both Ago1a and Ago1b, with Ago1a being the predominant form (**Fig. S7B**). To further explore the relative expression of Ago1a and Ago1b in aphid bacteriocytes, we analyzed publicly available RNAseq datasets from AphidBase (https://bipaa.genouest.org/is/aphidbase/), which integrates transcriptomic data from multiple studies for each aphid species. In parthenogenetic virginoparous females, Ago1a was predominantly expressed across all tissues, including bacteriocytes, whereas Ago1b expression was negligible in all tissues including bacteriocyte, relative to Ago1a (**Fig. S7C**).

Using this antibody, we then performed protein immunolocalization in stage 14-15 *A. pisum* embryos (**Fig. 2**). The experiment was independently repeated twice and included two controls: (i) a pre-adsorbed antibody control, where the antibody was incubated with purified recombinant Ago1a protein before tissue incubation to block binding to endogenous Ago1; and (ii) a secondary antibody-only control to exclude possible non-specific background fluorescence. Immunolocalization demonstrated that Ago1 is localized within *Buchnera* cells (**Fig. 2 and Video S1**) in *A. pisum* embryos. While Ago1 was ubiquitously detected throughout the embryos (**Fig. 2A-A^i^**), within bacteriocytes, the Ago1 signal was predominantly localized inside *Buchnera* cells, although some *Buchnera* cells lacked detectable signal (**Fig. 2A^ii^-A^iii^ and Video S1**). The within-*Buchnera* signal was largely depleted in the pre-adsorbed antibody control (**Fig. 2B-B^iii^**), while no background signal was detected in the secondary antibody-only control (**Fig. 2C-C^iii^**). We additionally performed the Ago1 immunolocalization experiment twice using the same antibody in *M. persicae*, and observed patterns of *Buchnera* localization (**Fig. S8**) consistent with those reported above in *A. pisum* (**Fig. 2**).

**Figure 2.**
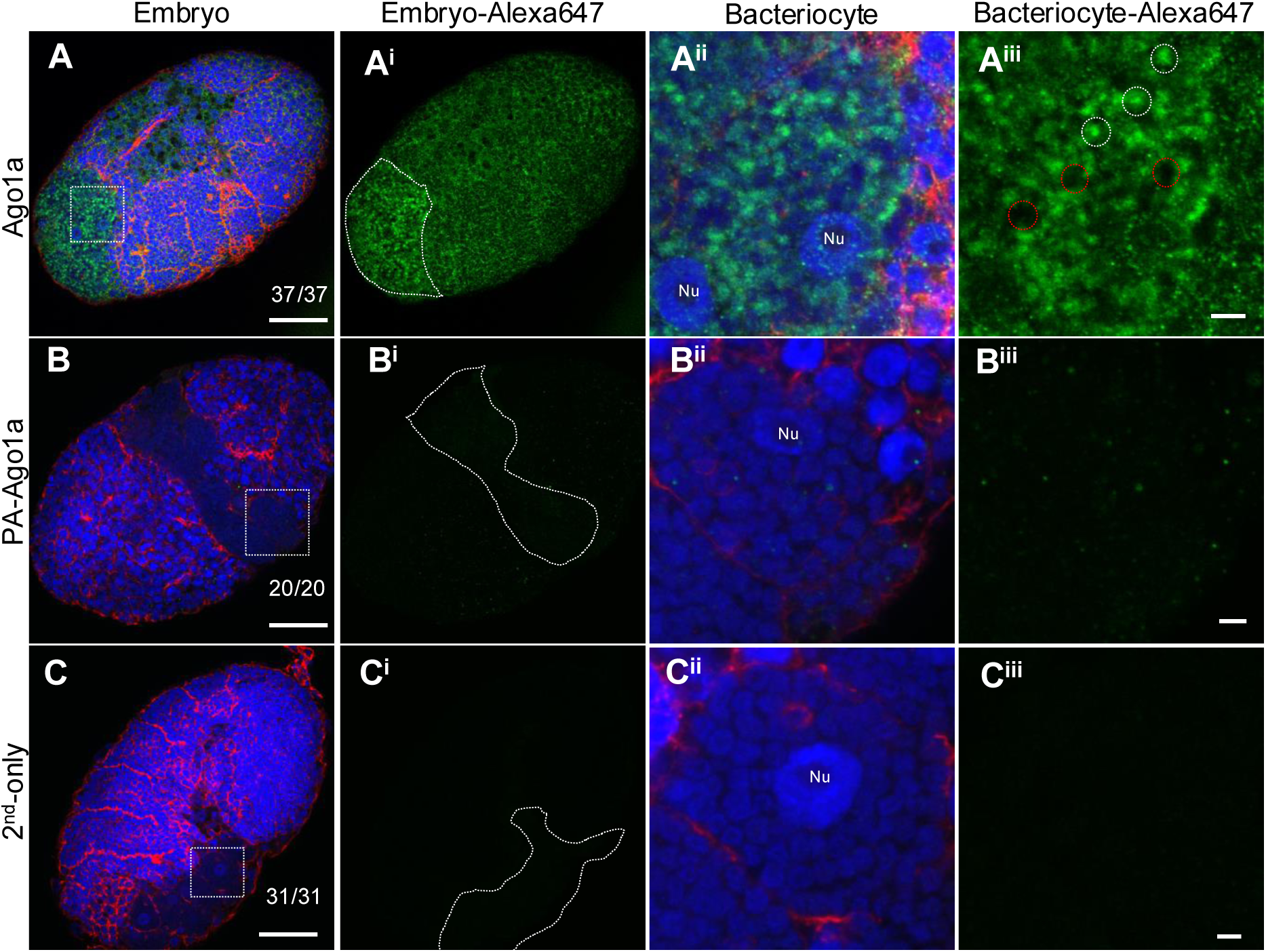
Ago1a immunolocalization in *A. pisum* embryos. All embryos are at stages 14-15 with clearly defined bacteriocytes. Blue indicates DAPI staining of DNA; Red indicates phalloidin staining of F-actin; Green indicates Ago1a localization. **(A–A^iii^)** Ago1a is ubiquitously distributed across embryos (A-A^i^). Within bacteriocytes, Ago1a signal is predominantly detected inside *Buchnera* cells (A^ii^-A^iii^). **(B–B^iii^)** The Ago1a signal is largely abolished in the pre-adsorbed control (PA-Ago1a). **(C– C^iii^)** No background signal is detected in secondary antibody-only control (2^nd^-only). Dashed squares in panel A-C mark regions shown at higher magnification in A^ii^-C^iii^. Bacteriome regions of the embryos are outlined in dashed lines in panels A^i^ - C^i^. Dashed circles in A^iii^ outline individual *Buchnera* cells; white cycles indicate *Buchenra* with Ago1a signal and red circles indicate those without Ago1a signal. Immunolocalization experiments were repeated twice for *A. pisum*. The number of embryos showing the observed patterns out of the total are summarized in the bottom right corner (A–D). All images were taken under the same confocal settings. Nu: bacteriocyte nucleus. Scale bars: 20 µm for embryo panels and 2 µm for bacteriocyte panels.

### Cross-kingdom translocated miRNAs are predicted to target Buchnera genes

Given that aphid miRNAs and Ago1a were localized within *Buchnera* cells, we predicted potential miRNA targets in the *Buchnera* genome following a eukaryotic miRNA target prediction framework. In this analysis, miRNA seed regions (nucleotides 2-8 at the 5’ end) were required to perfectly match target sequences, with additional matches in non-seed regions required to meet favorable binding energy constraints. Our predictions reveal that all four cross-kingdom translocated miRNAs (miR-1, miR-10, miR-29, and miR-927) have potential targets in *Buchnera* genes in both *A. pisum* and *M. persicae* (**Table S2A**). Conserved targets in *Buchnera* across the two host species were found for miR-1, miR-29, and miR-927, but not miR-10 (**Table S2B**). Notably, the conserved targets primarily represent four functional categories: (1) essential amino acid and vitamin biosynthesis including *carB* and *argH* for arginine biosynthesis, *dapF* for lysine biosynthesis, and *ribF* and *cysJ* for riboflavin biosynthesis; 2) components of the translational machinery, including tRNA synthesis for alanine, phenylalanine, and glutamine; and ribosomal proteins; 3) membrane-associated and structural proteins, including *flhA*, *fliK*, *flgI*, *ftsH*, and *murD* for flagellar assembly and peptidoglycan biosynthesis; 4) energy metabolism genes, including *atpA*, *clpX*, *hscA* and *pta* (**Table S2B**).

## Discussion

Using the aphid-*Buchnera* symbiosis as a model, we identified the cross-kingdom localization of four host encoded miRNAs within the cells of an ancient bacterial endosymbiont (**Fig. 1**). All four cross-kingdom translocated miRNAs have previously been implicated in host-microbe interactions or cross-kingdom regulation in other systems. Notably, in addition to localization within *Buchnera* cells in this study, miR-29 has also been found in aphid salivary glands and secreted into host plants during feeding (32), suggesting a broader role for this miRNA in inter-kingdom communication. While this is not direct evidence of cross-kingdom regulation, miR-927 has been shown to be upregulated in *Spodoptera exigua* cells in response to baculovirus infection (33). miR-10 is also broadly responsive to microbial challenge; from suppression in the nucleus of *Aedes aegypti* cell line infected with *Wolbachia* (34), and upregulation in *S. exigua* cells following baculovirus infection (33). Beyond insect-microbe systems, a homolog of miR-1 has been shown to be transported from the parasite, *Schistosoma japonicum*, into animal host cells where it targets the host gene grizzled, related protein 1 (*SFRP1*), promoting diseases progression (35). While we demonstrated that four are translocated to *Buchnera* cells, our results do not exclude the possibility that additional miRNAs undergo cross-kingdom translocation in aphids, for instance the bacteriome-abundant miRNAs (*e.g.,* miR-276 and miR-184a) and miRNAs that show higher, though not statistically significant, abundance in isolated *Buchnera* versus the bacteriome (*e.g.,* miR-2765 and let-7) (**Table S1**).

After showing that aphid miRNAs are translocated into *Buchnera* cells, we asked how these miRNAs might function within the symbiont. In insects, miRNA-mediated gene regulation requires loading of the mature miRNAs into the Ago1 protein to form the core RISC; because bacterial genomes lack Ago1, a key question is how cross-kingdom miRNAs might function in *Buchnera*. One possibility is that aphid miRNAs enter *Buchnera* together with host Ago1. Consistent with this hypothesis, our immunolocalization data showed that aphid Ago1 is specifically localized within *Buchnera* cells in both *A. pisum* and *M. persicae* (**Fig. 2**). The presence of Ago1 within *Buchnera* demonstrates that the essential target recognition module of the miRNA pathway is translocated into the symbiont. Whether additional components of the RISC complex are also transported, or whether Ago1-miRNA complexes regulate *Buchnera* gene expression through a non-canonical mechanism without additional host cofactors (26), remains to be determined. Nevertheless, our results support the possibility that translocated miRNAs may function in conjunction with host Ago1 to regulate *Buchnera* gene expression. Although the mechanism underlying this co-translocation remains unknown, host-to-symbiont protein trafficking is not unprecedented. For example, an aphid gene of bacterial origin encodes a protein that is transported into *Buchnera* (36), and aminotransferases encoded by the red palm weevil, *Rhynchophorus ferrugineus*, are imported into its endosymbiont, *Nardonella* to complete the tyrosine biosynthetic pathway (37).

Our data indeed suggest that aphid miRNAs function in *Buchnera* via Ago1a, however our results do not exclude alternative mechanisms. A pattern emerges in our data, at least for some miRNAs such that all *Buchnera* cells contain miRNA signal (**Fig. 1**), while some *Buchnera* cells lack detectable Ago1a signal (**Fig. 2A^ii^-A^iii^**). Thus, some cross-kingdom miRNAs may hijack endogenous gene-regulatory machinery of the endosymbiont. For example, in the whitefly *Bemisia tabaci,* miR-29 translocates into host tobacco plants and hijacks plant Ago1 to silence the defense gene *Bcl2-associated athanogene 4* (*Nt*BAG4) (32), and pathogen-derived miRNAs have been shown to rely on host Ago proteins for suppression of host immunity (38). In the absence of endogenous Agoa, the *Buchnera* genome encodes numerous conserved small RNAs that regulate bacterial gene expression (9) opening the possibility that aphid miRNAs may exploit *Buchnera*’s endogenous small RNA pathways.

The cross-kingdom translocation of Ago1a also provides insight into the form in which miRNAs are translocated. While multiple forms, such as precursor miRNAs, hairpins, and miRNA duplexes, have been shown to move across species boundaries, most documented cases involve single-stranded mature miRNAs being translocated (39, 40). Although we cannot fully exclude other forms, the presence of Ago1a (which binds mature miRNAs after duplex processing), detection of mature sequences in the *Buchnera* small RNAseq data, together with the specificity of our miRNA FISH probes for mature miRNA sequences, suggests that the miRNAs detected inside *Buchnera* are likely to be single-stranded mature miRNAs.

Following our determination that aphid miRNAs and Ago1a are translocated into *Buchnera* cells, we next set out to identify potential *Buchnera* genome targets. We found that all four cross-kingdom translocated miRNAs have putative targets in *Buchnera* (**Table S2A**). Interestingly, conserved *Buchnera* targets shared by *A. pisum* and *M. persicae* cluster into only four functional groups (**Table S2B**), including two that were previously demonstrated to be central to aphid-*Buchnera* symbiosis: essential amino acid and vitamin biosynthesis, and the membrane structural and transport machineries. Such a pattern suggests that the cross-kingdom translocated miRNAs are under positive selection regulating processes critical to symbiotic function. That said, these interactions are computational predictions that require functional validation. Experimental dissection of aphid miRNA and *Buchnera* target interactions *in vivo* remains challenging because of current technical limitations. However, recent advances provide potential avenues for such studies. For example, CRISPR-Cas9 genome editing has been developed to generate indel mutations using *A. pisum* sexual morphs (41, 42), offering a possible route to generate miRNA knockout lines. Even more promising, a newly developed *in vivo* method using synthetic single-stranded peptide nucleic acid (PNAs) to interfere with *Buchnera* gene expression (43), could be leveraged to test the functions of miRNAs specifically localized within *Buchnera* cells.

We have demonstrated the cross-kingdom translocation of aphid miRNAs and Ago1a into *Buchnera* cells, and yet the mechanism underlying this transport remains unresolved (**Fig. 3**). In other systems, cross-kingdom miRNA transfer occurs through several routes, including small RNA-binding proteins (39), high density lipoprotein complexes (44), cell-cell synapsis or gap-junctions (45), passive leakages due to cell injury or apoptosis (44), transmembrane proteins (45), as well as via exosomes or micro-extracellular vesicles (46). While all these pathways could be explored in aphids, transmembrane protein mediated transport and exosomes or extracellular vesicles trafficking are the most plausible mechanisms. First, for transmembrane proteins, systemic RNA interference deficient-1 (SID-1) is known to mediate the uptake of plant small RNAs including miRNAs into insects during feeding (47). Homologs of SID-1-like genes have been identified in the cotton/melon aphid, *Aphis gossypii*, and the grain aphid, *Sitobion avenae* using homology-based methods nearly two decades ago (48). A modern, genome-wide comparative approach is now needed to systematically annotate SID-1 homologs across aphid species. Identifying their expression patterns and localization, particularly within bacteriocytes, would help determine whether SID-1-mediated transport is a plausible mechanism for miRNA movement into *Buchnera*. In addition to SID-1, our previous immunolocalization work showed that the *A. pisum* non-essential amino acid transporter 1 is specifically localized to the symbiosomal membrane surrounding each *Buchnera* cell (49), supporting the possibility that aphid derived transmembrane proteins could contribute to this miRNA transfer. Conversely, although *Buchnera* possesses a highly reduced genome lacking flagellin genes, it retains proteins that assemble into hundreds of flagellar basal bodies on *Buchnera* cell surface (50). These flagellar basal bodies have been hypothesized to mediate material exchanges between aphid and *Buchnera* (51). In addition, several flagellar proteins were predicted to be targets of cross-kingdom miRNAs (**Table S2B**), raising the possibility of their involvement in miRNA translocation. Second, for exosomes or extracellular vesicles trafficking, electron microscopy has revealed that *Buchnera* cells are surrounded by numerous intracytoplasmic and intrasymbiosomal vesicles (52), and aphid bacteriocytes overexpress multiple vesicular trafficking pathways (53), together suggesting that exosomes or extracellular vesicles may also facilitate miRNA transport into *Buchnera*. Exosome or extracellular vesicle transport of miRNAs is often directed by sequence motifs in specific miRNAs (46). Among our candidates, only miR-1 carries a canonical 3’ 4-7 nucleotide EXOmotif (GGAG). MiR-29 and miR-10 possess core cell-retention-associated motif (CAGU for miR-29; GUAG for miR-10) (46), which may contribute to their accumulations in the bacteriocytes. However, no known EXOmotif is recognized in miR-10, miR-29 or miR-927, and the mechanisms that sort them into *Buchnera* remain to be elucidated.

**Figure 3.**
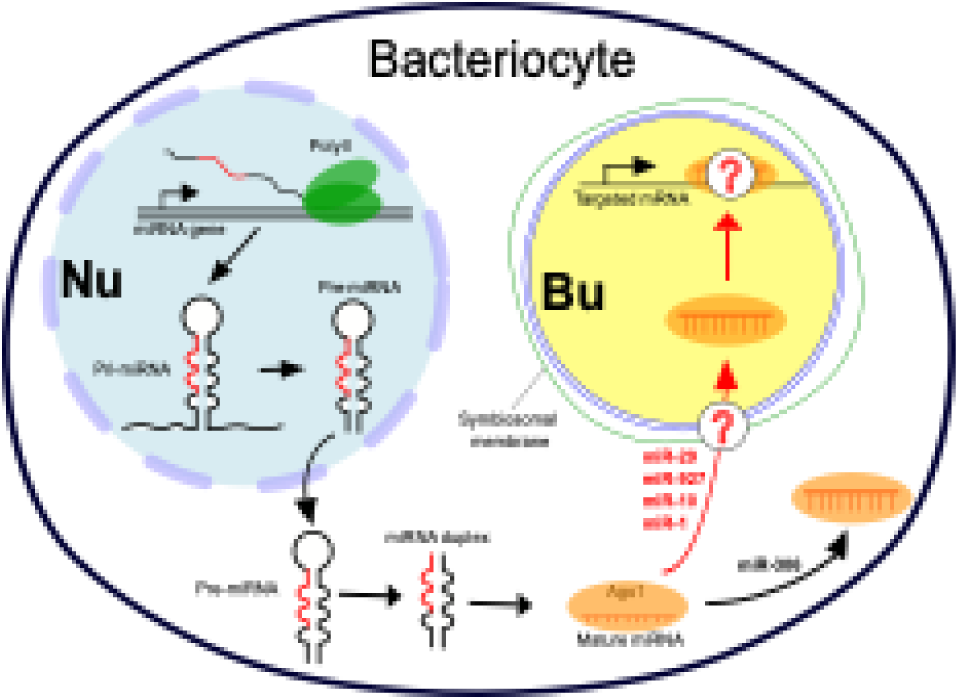
Synthesis of aphid miRNA cross-kingdom translocation into the endosymbiont *Buchnera*. In bacteriocyte nuclei, aphid miRNA genes are transcribed into primary miRNAs (pri-miRNAs), processed into precursor miRNAs (pre-miRNAs), and exported to the cytoplasm, where they are further processed into miRNA duplexes and loaded into the Argonaute protein as the core endonuclease. Mature miR-29, miR-927, miR-10, and miR-1 were detected within *Buchnera* cells, indicating cross-kingdom translocation, while miR-306 was not detected in *Buchnera* and appeared restricted to the bacteriocyte cytoplasm. The mechanism underlying miRNA endosymbiont translocation is unknown, including whether mature miRNAs move independently or in association with argonaute 1 (Ago1). Likewise, how these miRNAs may regulate gene expression within *Buchnera* remains unresolved. Nu: bacteriocyte nucleus; Bu: *Buchnera* cell; Yellow indicates detected aphid miRNA signals within *Buchnera*.

Our study raises many exciting questions about insect-endosymbiont communication. The identification of symbiosis-related miRNAs also highlights promising targets and tools for developing innovative pest control strategies. Given the indispensable nature of obligate nutritional endosymbiont to the survival of host insect pests, this symbiotic relationship has been explored as a vulnerability for pest control. For example, horizontally transferred genes such as *AmiD* and *LdcA1*, which are essential for maintaining the aphid-*Buchnera* symbiosis, have been successfully targeted by RNA interference. Knockdown of these genes significantly reduces *Buchnera* abundance and activity, leading to impaired aphid growth and performance (54). miRNA-based approaches are also emerging as powerful platforms for pest management. Artificial miRNAs (amiRNAs) have been engineered for expression in host plants to control pests like the cotton bollworm, *Helicoverpa armigera* (55) and *M. persicae* (56, 57). In addition, aphid miR-3024 recently was shown to modulate interactions between aphid and the secondary symbiont *Serratia symbiotica*. Overexpression of miR-3024 in host plants suppressed the aphid *MRP4* gene, a transporter of vitamin B6 (VB6) supplied by *S. symbiotica*, ultimately causing aphid mortality (15). Notably, miR-3024 and miRNA-treated aphids did not adversely affect beneficial insects like ladybeetles and bumble bees (15). Our discovery that aphid-derived miRNAs localize within its obligate symbiont *Buchnera* identifies a new class of regulatory molecules that could be leveraged as precise and potential targets for aphid control.

In conclusion, we have shown that aphid-derived miRNAs localize to *Buchnera* cells, revealing a previously unknown layer of communication in an ancient insect-bacterial endosymbiosis. Our findings provide strong evidence for cross-kingdom miRNA transport. Given the essential nature of *Buchnera* to aphids and the agricultural importance of aphids as pests, these symbiosis-related miRNAs offer compelling opportunities for innovative, miRNA- and symbiosis-based pest management strategies.

## Materials and Methods

### Aphid cultures

The aphid line, *Acyrthosiphon pisum* LSR1 (58), was maintained on 4-5 true leaf stage broad beans (*Vicia faba)* (Todd’s seeds®) using plastic cages (30 x 30 x 30 cm^3^). The aphid line, *Myzus persica*e USDA-Red (59, 60), was maintained on *Nicotiana tabacum* wild-type leaves using plastic cages (40 x 40 x 60 cm^3^). Broad bean plants and wild type *Nicotiana tabacum* plants were grown in commercial soil Miracle-Gro® Potting Mix with Osmocote Plus™ fertilizer (∼5 g were applied manually around each plant). The aphid colony and plants were maintained in a climate-controlled Conviron GEN 1000 growth chamber with the following settings: 23°C, with a 16:8 hour light:dark cycle (100% LED-W) in the laboratory.

### Differentially expressed miRNAs identification by small RNAseq

We mined *Buchnera* small RNAseq data generated by Hansen and Degnan (9) for aphid miRNAs. The samples included *Buchnera* strains from four aphid species: *A. pisum* (*Buchnera*-LSR1 and *Buchnera*-5A), *A. kondoi* (*Buchnera*-Ak), *Uroleucon ambrosiae* UA002 (*Buchnera*-Ua), and *Schizaphis graminum* (*Buchnera*-Sg). The raw data were downloaded from NCBI under the accession number: PRJNA212118 (5A-SRR935066, LSR1-SRR935070, Ak-SRR935071, Ua-SRR935072 and Sg-SRR935073).

To identify aphid miRNAs enriched in *Buchnera*, the raw reads from *Buchnera* small RNAseq data were mapped to our reference aphid miRNAs (generated from multiple lines of *A. pisum* and *M. persicae*) using Bowtie with the same settings (20). The read counts for each miRNAs detected in the *Buchnera* small RNAseq datasets were compiled together with their corresponding counts from the bacteriome RNAseq datasets (20) into a count matrix for differential expression analysis. Principal component analysis and differentially expression analyses were conducted using Bioconductor package edgeR v.3.10.2 (61) with each of the *Buchnera* and aphid lines treated as biological replicate. miRNAs with *p* <= 0.05 and FDR <= 0.05 were retained as differentially expressed.

### miRNA fluorescence in situ hybridization

We performed miRNA fluorescence *in situ* hybridization (FISH) on aphid developing embryos, focusing on stages after bacteriocytes cells form (stage 14-15). Embryos were dissected from 10-15 ∼3-day old aphid nymphs and placed in 1 x PBS (VWR, USA). Samples were washed three times using 1 x PBS to remove fat bodies and fixed in 4% formaldehyde in methanol overnight at 4 °C. Following fixation, samples were permeabilized with 0.1% Triton X-100 for 10 minutes and treated with 0.075% (vol/vol) Proteinase K in buffer (5 mM Tris-HCl, 1 mM EDTA, 1 mM NaCl in RNase-free water) for 10 minutes at 37 °C to release protein-bound RNAs. Hybridization then was performed overnight at probe-specific temperatures using 5’TYE665-labeled, miRNA-specific miRCURY LNA miRNA Detection Probes (Qiagen, US, **Table S3**) in miRNA ISH buffer (Qiagen, US). The hybridization temperatures were calculated as 20 °C below the TM of each miRNA probe. After hybridization, samples were washed sequentially with 5x, 2.5x, and 1.25x saline-sodium citrate buffer (SSC) (Quality Biological, USA). Genomic DNA was counterstained with DAPI (1:1000) (Invitrogen, US), and F-actin was stained with Phalloidin (1:20) (Biotium, US) to facilitate tissue visualization. Finally, samples were mounted to slides using ProLong Diamond Antifade Mountant (Thermo Fisher Scientific, USA) and imaged under a Leica SP8 confocal microscope located at the Advanced Microscopy and Analytical Core (AMAC) at Louisiana State University. Between each step, samples were washed three times using 1x PBST before the probe hybridization step and 1.25x SSC after hybridization. To ensure probe specificity and protocol integrity, positive controls were performed with probes targeting the small nucleus RNA U6. Negative controls included a preabsorbed probe treatment, where miRNA probes were incubated overnight at hybridization temperature with a 4-fold concentration of corresponding miRNA mimics, and a no-probe control to exclude non-specific and background fluorescence. All miRNA detection experiments and controls were independently repeated 2-3 times for both *A. pisum* and *M. persicae*.

To quantify fluorescent signal intensity around *Buchnera* cells, we performed line-scan intensity measurements across individual *Buchnera*. All measurements were conducted on images that were maginified to the level of single bacteriocyte cells. For each image, random measurement points were generated using Fiji macro within regions displaying clear and circular *Buchnera* identified by DAPI staining (ImageJ 1.54P). A line was drawn across each selected *Buchnera* to extract signal intensity profiles along the line. For each probe, we measured five randomly selected *Buchnera* per image, and a total of over 10 images across three independent experiments (over 50 measurements in total per treatment). To integrate data across samples, *Buchnera* cell membranes were used as positional boundaries and line positions were normalized accordingly. At each normalized position, mean fluorescence intensities for both DAPI and TYE665 (for miRNA), along with standard errors, were calculated and plotted in RStudio v2024.12.0+467.

### Argonaute 1 protein immunolocalization

For antibody production, the full-length *A. pisum* Argonaute 1 coding sequence was submitted to GenScript. A mouse-derived polyclonal antibody was generated against the entire Ago1a protein. The generated antibody was first validated through Western blot using whole-aphid lysates with the purified protein as positive control (**Fig. S7B**). The protein bands corresponding to the immunoreactive signals detected by Western blot were excised from an SDS-PAGE gel and subjected to mass spectrometry analysis to confirm their identifies. This antibody was used for immunolocalization experiments to test the potential cross-kingdom translocation of Ago1a from aphid to *Buchnera*.

For protein immunolocalization, aphid embryos were dissected from 10 to 15 young adult females in 1x PBS (VWR, USA) and fixed overnight at 4 °C in 4% (wt/vol) formaldehyde (Thermo Scientific) prepared in 1x PBS, and then washed 3× (15 min per wash) in 1x PBS at room temperature. Permeabilization was conducted with 0.1% (vol/vol) Triton X-100 for 10 min washed 3× (15 min per wash) in PBS with 0.05% (vol/vol) Tween (PBST). Embryos were then treated with 0.05% (vol/vol) proteinase K for 10 min at 37°C and blocked with 3% (vol/vol) bovine serum albumin (BSA) in PBST for 1 h at room temperature. Samples were then incubated with primary anti-Ago1a antibody 1:250 in 1% BSA in PBST overnight at 4 °C. The next day, embryos were washed and incubated with Alexa-Fluor^TM^ 647 goat anti-mouse IgG (H+L) secondary antibody (Thermo Fisher Scientific, USA) 1:250 in 1% BSA in PBST 1 hours at room temperature. Embryos were stained with DAPI (1:1000) (Invitrogen, US), and phalloidin (1:20) (Biotium, US) for 30 min at room temperature. After incubation, samples were mixed with ProLong Diamond Antifade Mountant (Thermo Fisher Scientific, USA) and mounted in 2,2′-thiodiethanol (Sigma-Aldrich) on a glass slide. Slides were imaged using a Leica SP8 confocal microscope located at the AMAC core at LSU. Controls were run in parallel and included localizations with purified protein-preabsorbed primary antibody (using a 10-fold molar excess of purified protein) and localizations with the secondary antibody only. The localization experiment, with control treatments, was repeated twice with both *A. pisum* and *M. persicae* samples. In each experiment, multiple individual embryos at stage 14-15 were imaged, and z-stack images were captured to further confirm the translocation of signals in three dimensions (**Video S1**).

### miRNA target prediction in Buchnera

We predicted the potential target sites of four miRNAs that are found inside of *Buchnera* using the coding sequences of mRNAs for both *Buchnera*Mp str.USDA (GenBank#: GCA_000521525.1) and *Buchnera*Ap str.LSR1 (GenBank#:GCA000174075.1) using miRanda v.3.3a (Enright et al., 2004). Potential target sites were predicted and filtered using the following parameters: -sc 80 -go -8 -ge -2 -scale 2 -en 0 -strict -quiet, where we demanded no mismatches in the 5’ seed region (6 nt in length) (**Table S2A**). For all predicted miRNA and mRNA pairs, we further extracted the common ones across *Buchnera*Mp Str.USDA and *Buchnera*Ap str.LSR1. We annotated these common targets by assigning them to pathways using the Kyoto Encyclopedia of Genes and Genomes (KEGG: http://www.kegg.jp) and specific functions through UniPort (https://www.uniprot.org) (**Table S2B**).

## Data, Materials, and Software Availability

Supplemental tables, figures and videos are provided in *supporting information*.

## Acknowledgement

We thank Dr. Hsiao-ling Lu from National Formosa University, Taiwan, for her valuable suggestions on selecting miRNA *in situ* hybridization probes to avoid *Buchnera* autofluorescence. We thank Dr. Anastasios Vourekas and Josef Mick, from the Department of Biology, Louisiana State University, for their advice on Argonaute protein antibody generation and preliminary western blot tests. This research was supported by start-up funds from the Department of Entomology, Louisiana State University, to H.F.; National Science Foundation Awards IOS-1121847 and IOS-1354154 to A.C.C.W.; and United States Department of Agriculture - Agriculture and Food Research Initiative award 2021-67013-33565 to G.J.

## Author contributions

HF, ACCW and HS designed research; YD, HS, and HF performed research; YD, GJ, ACCW, and HF analyzed data; YD and HF drafted the manuscript; GJ, ACCW, and HF revised the manuscript; GJ, ACCW, and HF provided funding.

---

**Figure S1.**
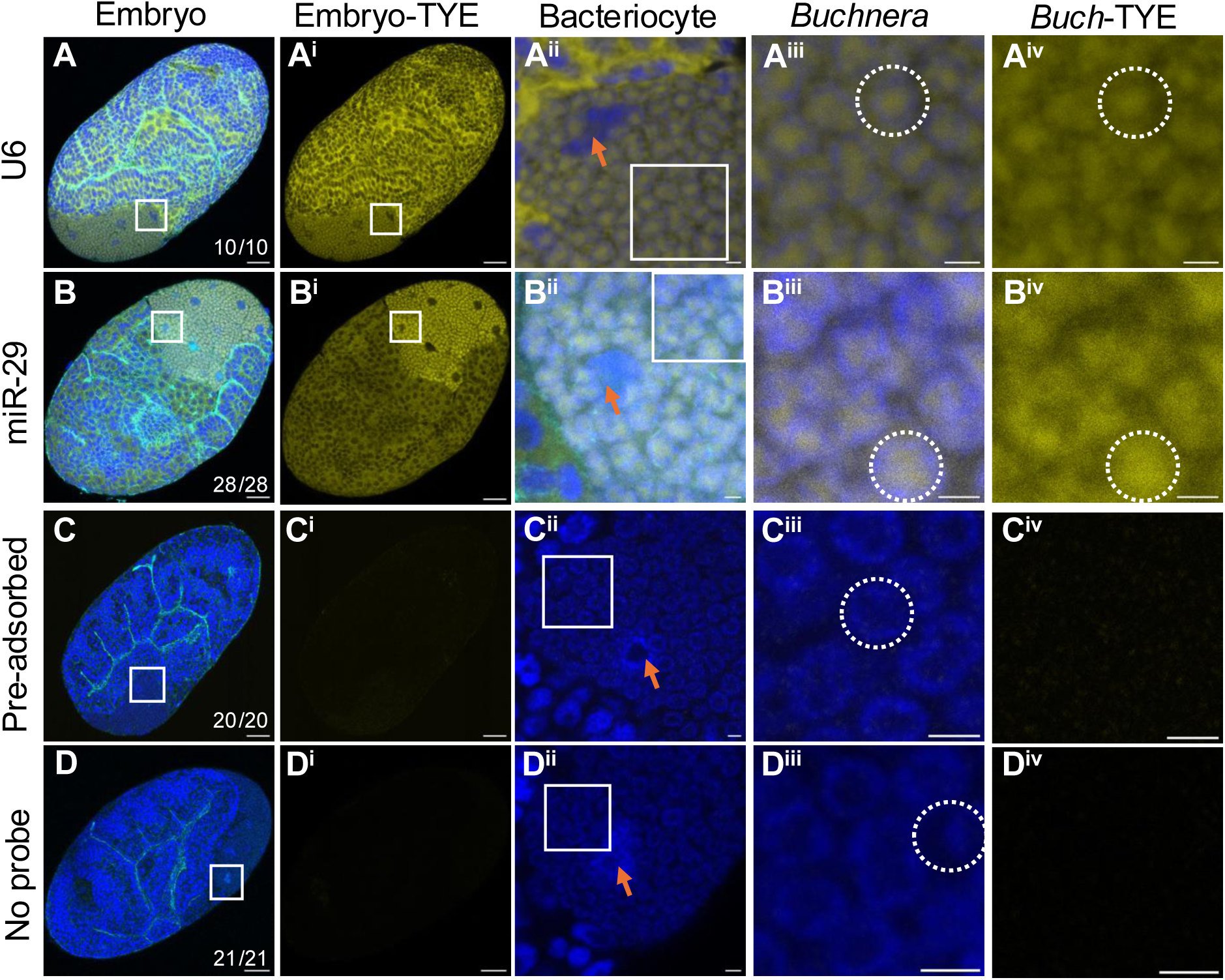
Complete experimental design for miR-29 fluorescent *in situ* hybridization (images from *M. persicae* samples). Blue indicates DAPI staining of DNA; Cyan indicates phalloidin staining of F-actin; Yellow indicates miRNA localization visualized using the TYE665 (TYE) fluorophore. Squares indicate regions shown at higher magnification in the adjacent columns. **(A–A^iv^)** U6, a ubiquitous small nuclear RNA, is enriched in the nucleolus of all aphid cells (A-A^iii^), and its signal in bacteriocyte is localized to the bacteriocyte nucleus (A^ii^) and cytoplasm (A^iv^). **(B–B^iv^)** miR-29 is significantly enriched in bacteriocytes (B^i^ & C^i^). Higher magnification (B^ii^-B^iv^) reveals that miR-29 localizes within *Buchnera* cells (dashed circles). No specific signals were detected in pre-adsorbed (**C-C^iv^**) and no-probe controls (**D-D^iv^**) across all experiments. The complete FISH experiment was repeated three times across *A. pisum* and *M. persicae*. The number of embryos showing the observed patterns out of the total are summarized in the bottom right corner (A–D). All images were taken under the same confocal settings. Scale bars represent 20 µm in panels with embryos, and 2 µm in panels with bacteriocytes or *Buchnera* cells.

**Figure S2.**
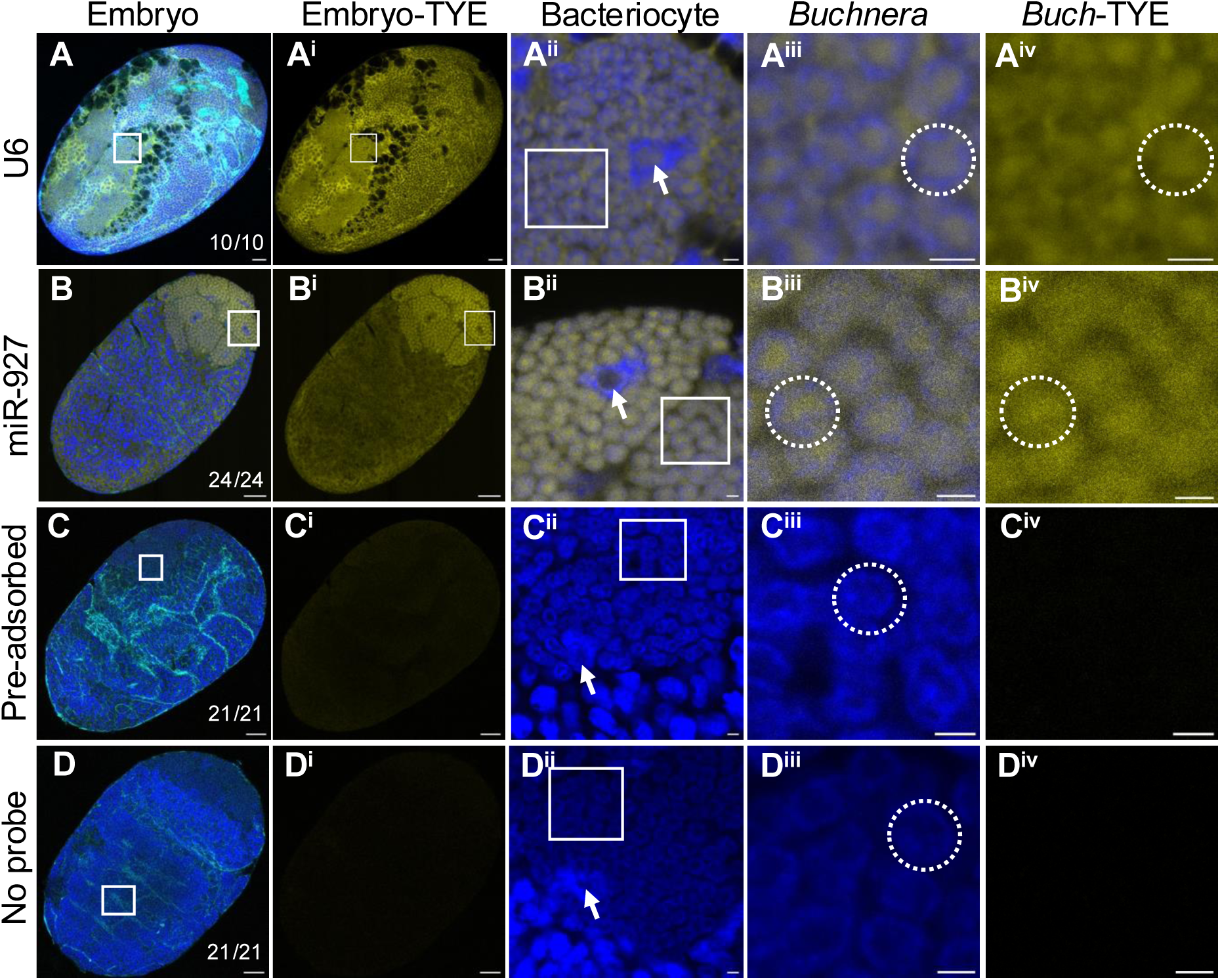
Complete experimental design for miR-927 fluorescent *in situ* hybridization (images from *M. persicae* samples). Blue indicates DAPI staining of DNA; Cyan indicates phalloidin staining of F-actin; Yellow indicates miRNA localization visualized using the TYE665 (TYE) fluorophore. Squares indicate regions shown at higher magnification in the adjacent columns. **(A–A^iv^)** U6, a ubiquitous small nuclear RNA, is enriched in the nucleolus of all aphid cells (A-A^iii^), and its signal in bacteriocyte is localized to the bacteriocyte nucleus (A^ii^) and cytoplasm (A^iv^). **(B–B^iv^)** miR-927 is significantly enriched in bacteriocytes (B^i^ & C^i^). Higher magnification (B^ii^-B^iv^) reveals that miR-927 localizes within *Buchnera* cells as shown in puncta (dashed circles). No specific signals were detected in pre-adsorbed (**C-C^iv^**) and no-probe controls (**D-D^iv^**) across all experiments. The complete FISH experiment was repeated three times across *A. pisum* and *M. persicae*. The number of embryos showing the observed patterns out of the total are summarized in the bottom right corner (A–D). All images were taken under the same confocal settings. Scale bars represent 20 µm in panels with embryos, and 2 µm in panels with bacteriocytes or *Buchnera* cells.

**Figure S3.**
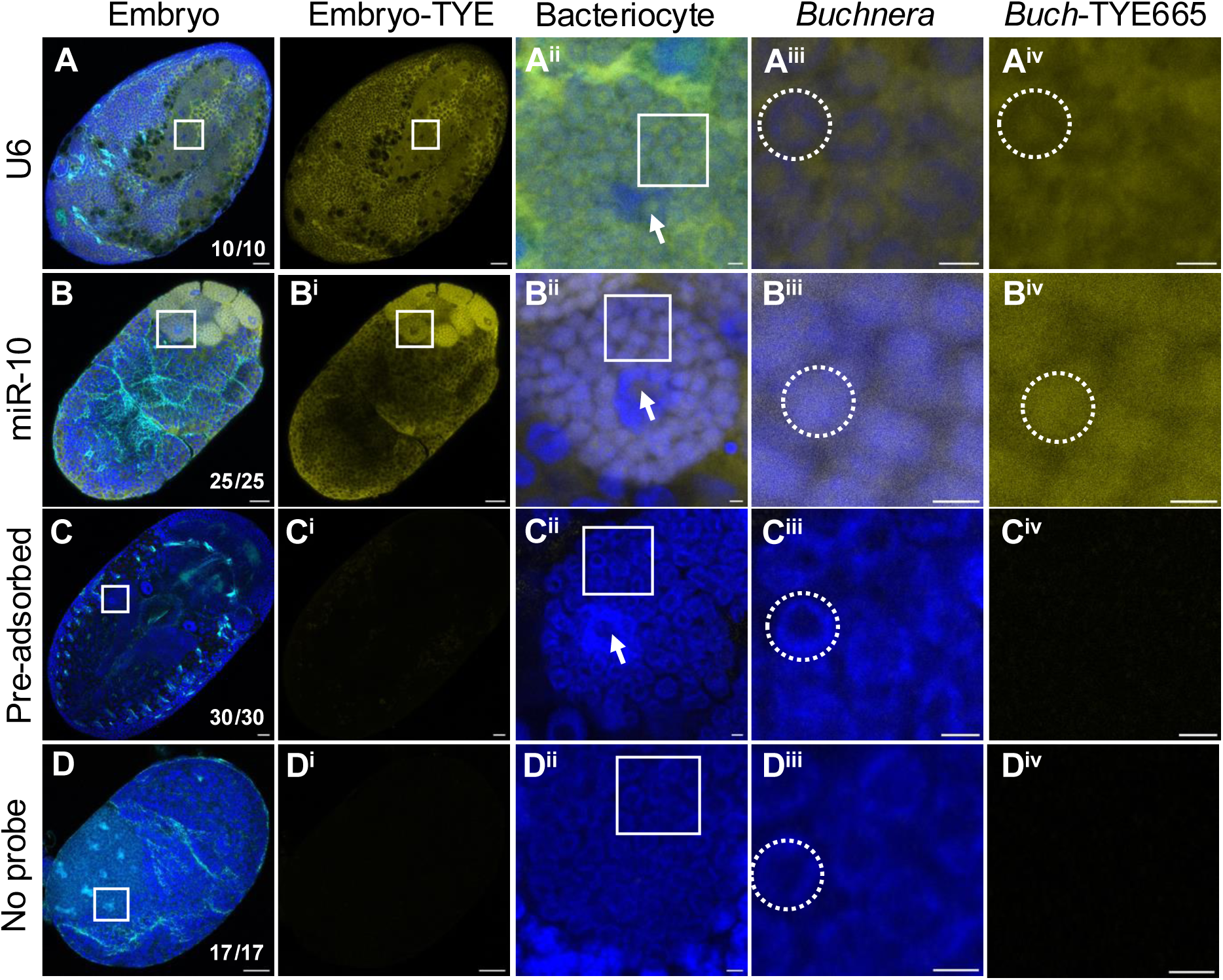
Complete experimental design for miR-10 fluorescent *in situ* hybridization (images from *M. persicae* samples). Blue indicates DAPI staining of DNA; Cyan indicates phalloidin staining of F-actin; Yellow indicates miRNA localization visualized using the TYE665 fluorophore. Squares indicate regions shown at higher magnification in the adjacent columns. **(A–A^iv^)** U6, a ubiquitous small nuclear RNA, is enriched in the nucleolus of all aphid cells (A-A^iii^), and its signal in bacteriocyte is localized to the bacteriocyte nucleus (A^ii^) and cytoplasm (A^iv^). **(B–B^iv^)** miR-10 is ubiquitously expressed across the embryos including bacteriocytes (B^i^ & C^i^). Higher magnification (B^ii^-B^iv^) reveals that miR-10 localizes within *Buchnera* cells without clear *Buchnera* cell boundaries (dashed circles). No specific signals were detected in pre-adsorbed (**C-C^iv^**) and no-probe controls (**D-D^iv^**) across all experiments. The complete FISH experiment was repeated three times across *A. pisum* and *M. persicae*. The number of embryos showing the observed patterns out of the total are summarized in the bottom right corner (A–D). All images were taken under the same confocal settings. Scale bars represent 20 µm in panels with embryos, and 2 µm in panels with bacteriocytes or *Buchnera* cells.

**Figure S4.**
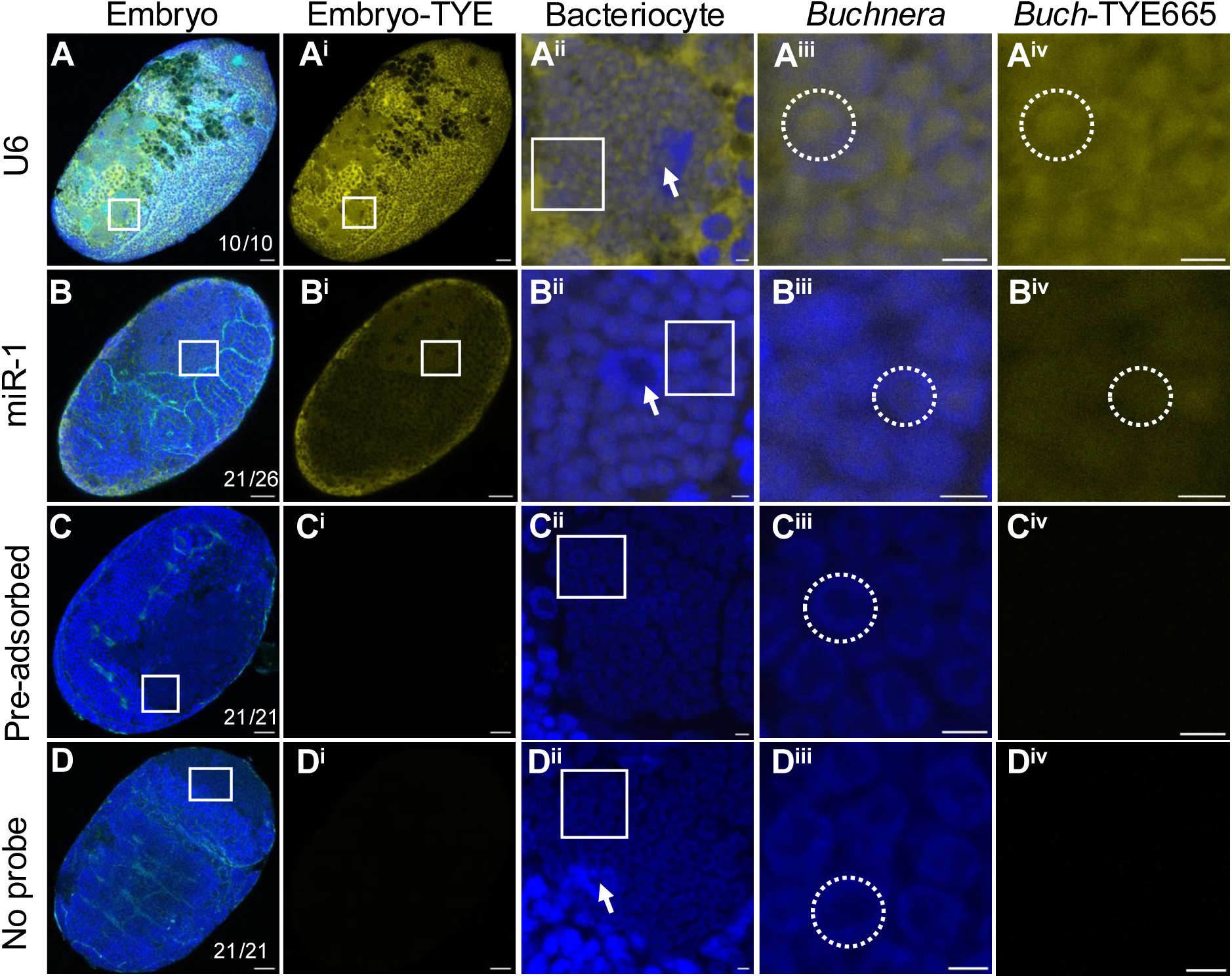
Complete experimental design for miR-1 fluorescent *in situ* hybridization (images from *M. persicae* samples). Blue indicates DAPI staining of DNA; Cyan indicates phalloidin staining of F-actin; Yellow indicates miRNA localization visualized using the TYE665 fluorophore. Squares indicate regions shown at higher magnification in the adjacent columns. **(A–A^iv^)** U6, a ubiquitous small nuclear RNA, is enriched in the nucleolus of all aphid cells (A-A^iii^), and its signal in bacteriocyte is localized to the bacteriocyte nucleus (A^ii^) and cytoplasm (A^iv^). **(B–B^iv^)** miR-1 is lowly detected in bacteriocytes (B^i^ & C^i^). Higher magnification (B^ii^-B^iv^) reveals that miR-1 signals are likely localized within *Buchnera* cells (dashed circles). No specific signals were detected in pre-adsorbed (**C-C^iv^**) and no-probe controls (**D-D^iv^**) across all experiments. The complete FISH experiment was repeated three times across *A. pisum* and *M. persicae*. The number of embryos showing the observed patterns out of the total are summarized in the bottom right corner (A–D). All images were taken under the same confocal settings. Scale bars represent 20 µm in panels with embryos, and 2 µm in panels with bacteriocytes or *Buchnera* cells.

**Figure S5.**
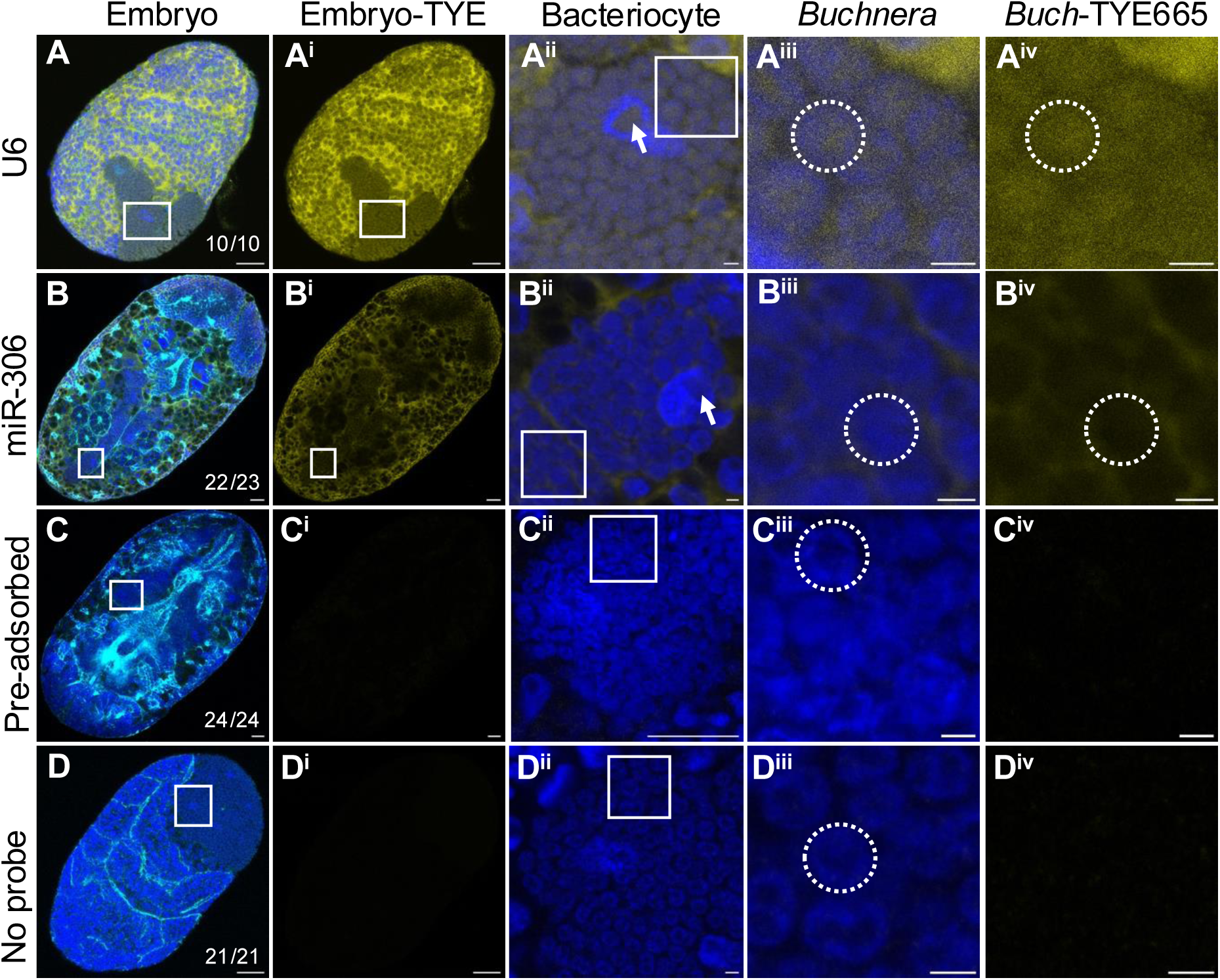
Complete experimental design for miR-306 fluorescent *in situ* hybridization (images from *M. persicae* samples). Blue indicates DAPI staining of DNA; Cyan indicates phalloidin staining of F-actin; Yellow indicates miRNA localization visualized using the TYE665 fluorophore. Squares indicate regions shown at higher magnification in the adjacent columns. **(A–A^iv^)** U6, a ubiquitous small nuclear RNA, is enriched in the nucleolus of all aphid cells (A-A^iii^), and its signal in bacteriocytes is localized to the bacteriocyte nucleus (A^ii^) and cytoplasm (A^iv^). **(B–B^iv^)** miR-306 is detected in bacteriocytes, but not as high abundance as other tissues (B-B^ii^). Higher magnification (B^ii^-B^iv^) reveals that miR-306 localizes in the cytoplasm and not within *Buchnera* cells (dashed circles). No specific signals were detected in pre-adsorbed (**C-C^iv^**) and no-probe controls (**D-D^iv^**) across all experiments. The complete FISH experiment was repeated three times across *A. pisum* and *M. persicae*. The number of embryos showing the observed patterns out of the total are summarized in the bottom right corner (A–D). All images were taken under the same confocal settings. Scale bars represent 20 µm in panels with embryos, and 2 µm in panels with bacteriocytes or *Buchnera* cells.

**Figure S6.**
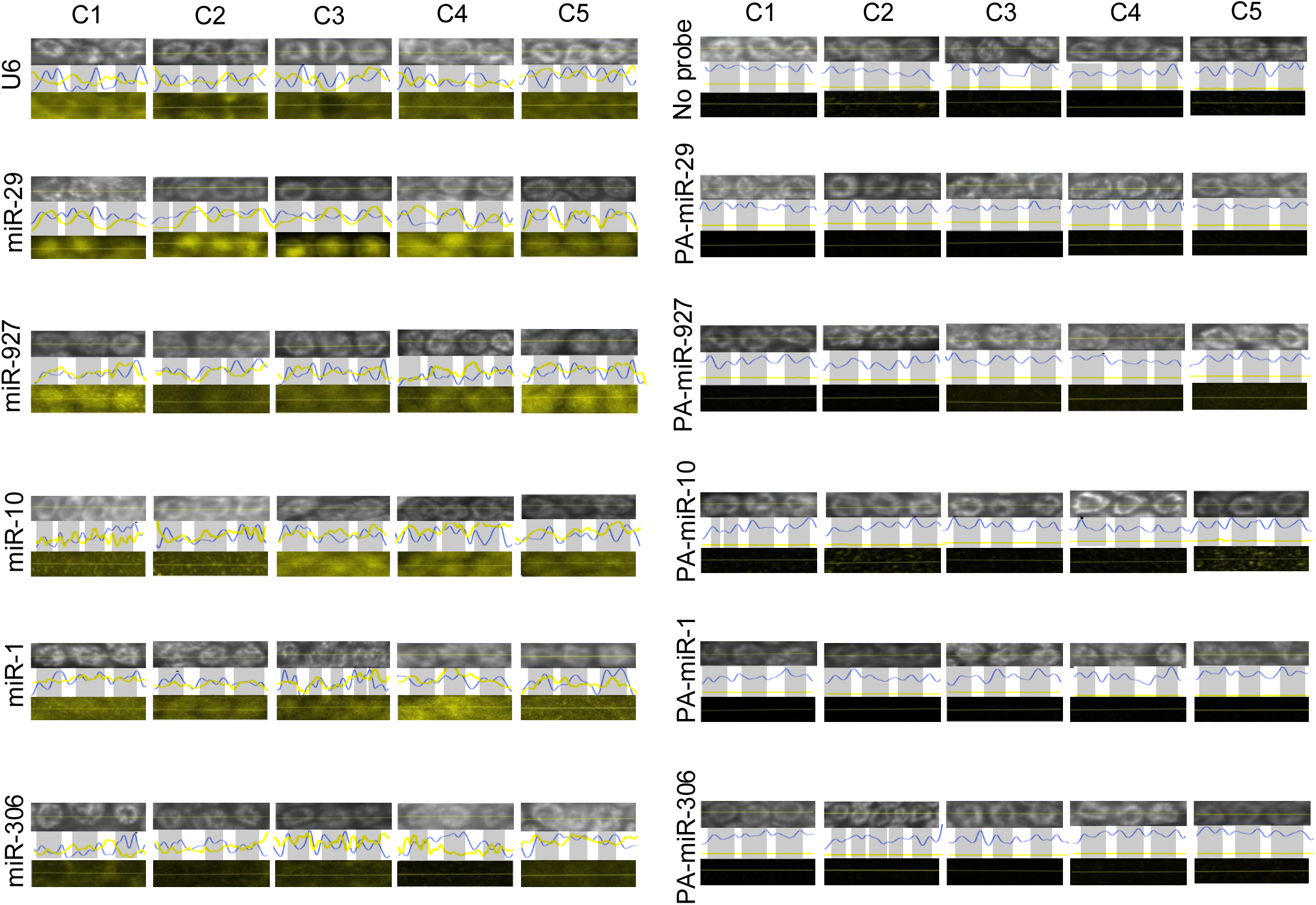
Line-scan intensities across multiple *Buchnera* cell spaces. For each miRNA, five bacteriocyte cells (represented as individual blocks labeled as C1-C5) were randomly selected from different embryos across independent experiments. Within each bacteriocyte, line-scan was performed across at least three *Buchnera* cells. Each block consists of three horizontal panels: the top panel shows DAPI staining, the middle panel displays the corresponding line-scan intensity profiles for DAPI (blue, shaded blocks indicate *Buchnera* boundaries) and TYE665 (yellow), and the bottom panel shows the TYE665 fluorescence of the miRNA within the same scanned region. The No-probe control includes five bacteriocyte cells, each randomly selected from the corresponding control experiment for each miRNA.

**Figure S7.**
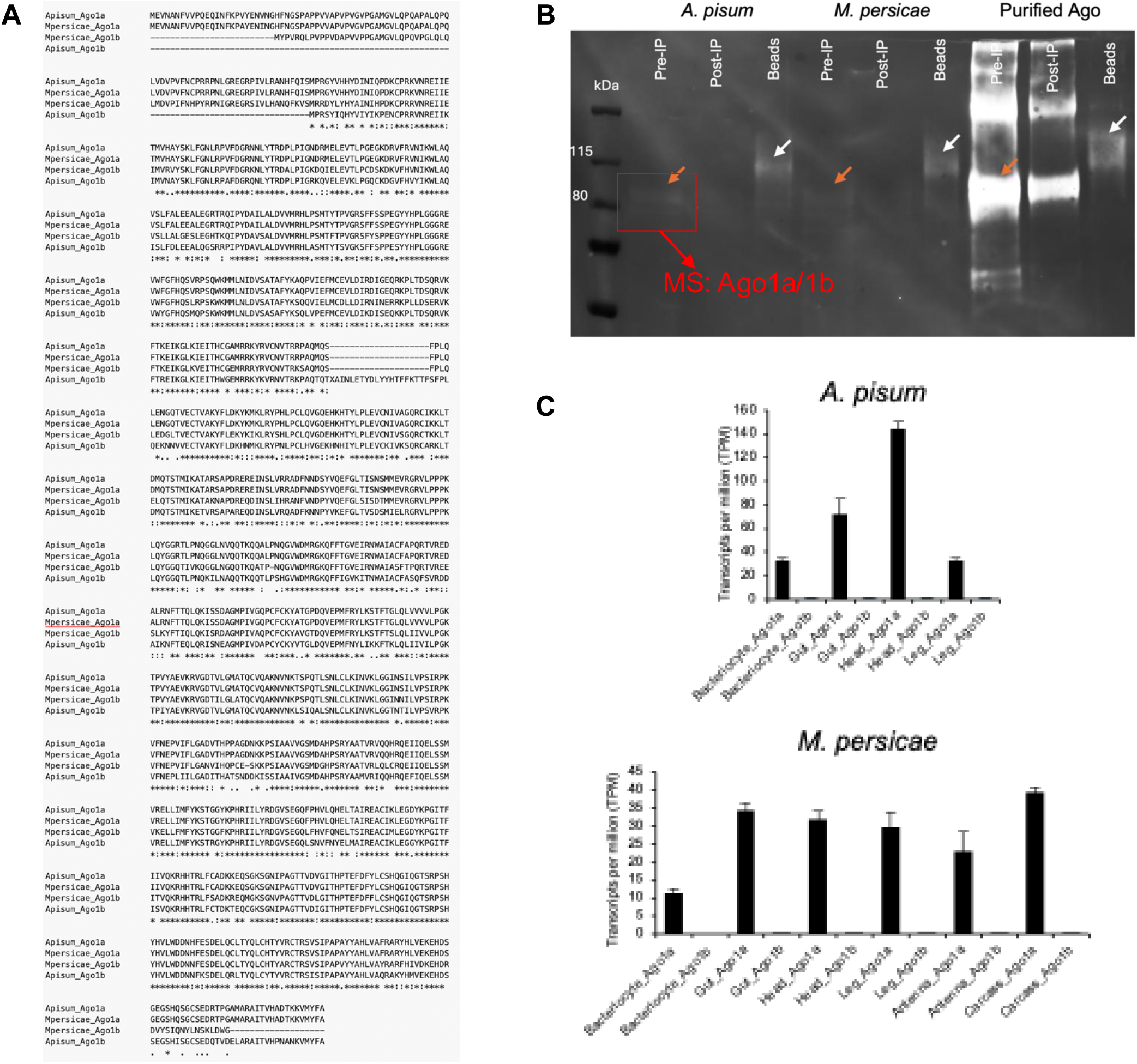
Western blot confirmation of Ago1a antibody specificity in *A. pisum* and *M. persicae*. (**A**) Sequence alignment of Ago1a and Ago1b from *A. pisum* and *M. persicae*. (**B**) Western blot analysis of protein extracts from the two aphid species (300 mg of whole aphids per species) using the Ago1a antibody, with purified Ago1a protein as a positive control. For each sample (*A. pisum*, *M. persicae,* and purified Ago), three fractions were analyzed : 1) Pre-IP: total protein before immunoprecipitation (IP) with Ago1a antibody-coated beads; 2) Post-IP: the unbound protein fraction after IP; 3) Beads: proteins eluted from the antibody-coated beads. Orange arrows indicate the specific bands in the Pre-IP samples corresponding to the expected size of Ago1 proteins, which were excised for mass spectrometry (MS) analysis. MS detected both Ago1a and Ago1b peptides, with Ago1a representing the predominant protein. White arrows indicate proteins eluted from the IP beads. The bands shifted slightly in size but remained consistent with the purified Ago control. These specific bands were largely depleted in the Post-IP fractions of both aphid samples and markedly reduced in the purified Ago1a control, suggesting that the antibody specifically recognizes and enriches Ago1a proteins. (**C**) Tissue-specific expression of Ago1a and Ago1b in *A. pisum* LSR1 (n=18 for bacteriocyte, n=7 for gut, and n=6 for head and leg) and *M. persicae* clones 5410R and NS (n=4 for all tissues). Expression data were retrieved from from AphidBase and compliled from multiple RNAseq datasets, primarily generated from parthenogenetic female aphids. Ago1b expression was negligible across all tissues, including bacteriocytes, compared with Ago1a.

**Figure S8.**
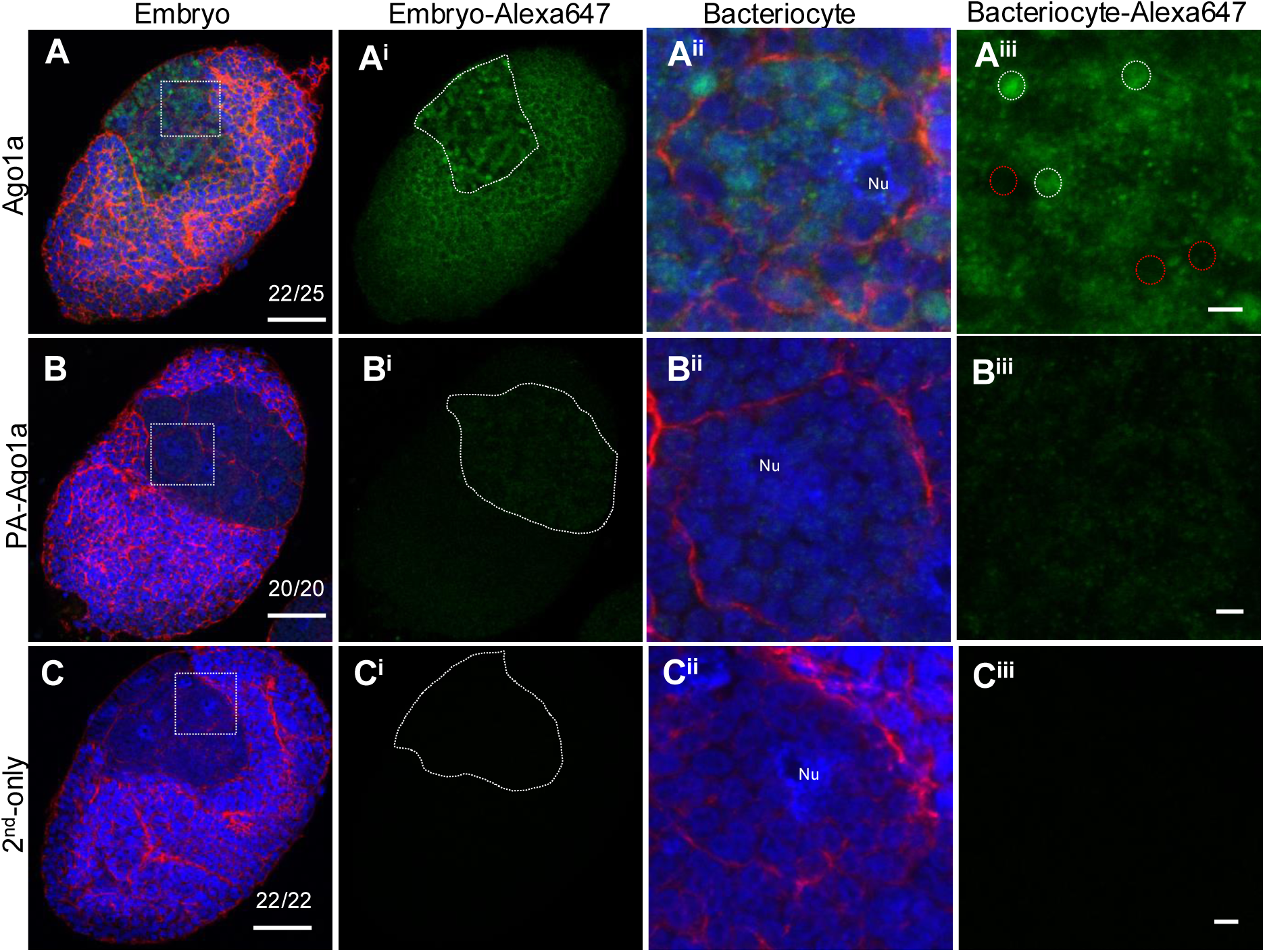
Ago1 immunolocalization in *M. persicae* embryos. All embryos are at stages 14-15 with clearly defined bacteriocytes. Blue indicates DAPI staining of DNA; Red indicates phalloidin staining of F-actin; Green indicates Ago1a localization. **(A–A^iii^)** Ago1a is ubiquitously distributed across embryos (A-A^i^). Within bacteriocytes, Ago1a signal is predominantly detected inside *Buchnera* cells (A^ii^ - A^iii^). **(B–B^iii^)** The Ago1a signal is largely abolished in the pre-adsorbed control (PA-Ago1a). **(C–C^iii^)** No background signal is detected in secondary antibody-only control (2^nd^-only). Dashed squares in panel A-C mark regions shown at higher magnification in A^ii^-C^iii^. Bacteriome regions of the embryos were outlined in dashed lines in panels A^i^ - C^i^. Dashed circles in A^iii^ outline individual *Buchnera* cells; white cycles indicate *Buchnera* with Ago1a signal and red circles indicate those without Ago1a signal. Immunolocalization experiments were repeated twice for *M. persicae*. The number of embryos showing the observed patterns out of the total are summarized in the bottom right corner (A–D). All images were taken under the same confocal settings. Nu: bacteriocyte nucleus. Scale bars: 20 µm for embryo panels and 2 µm for bacteriocyte panels.

**Supplemental Video 1:**
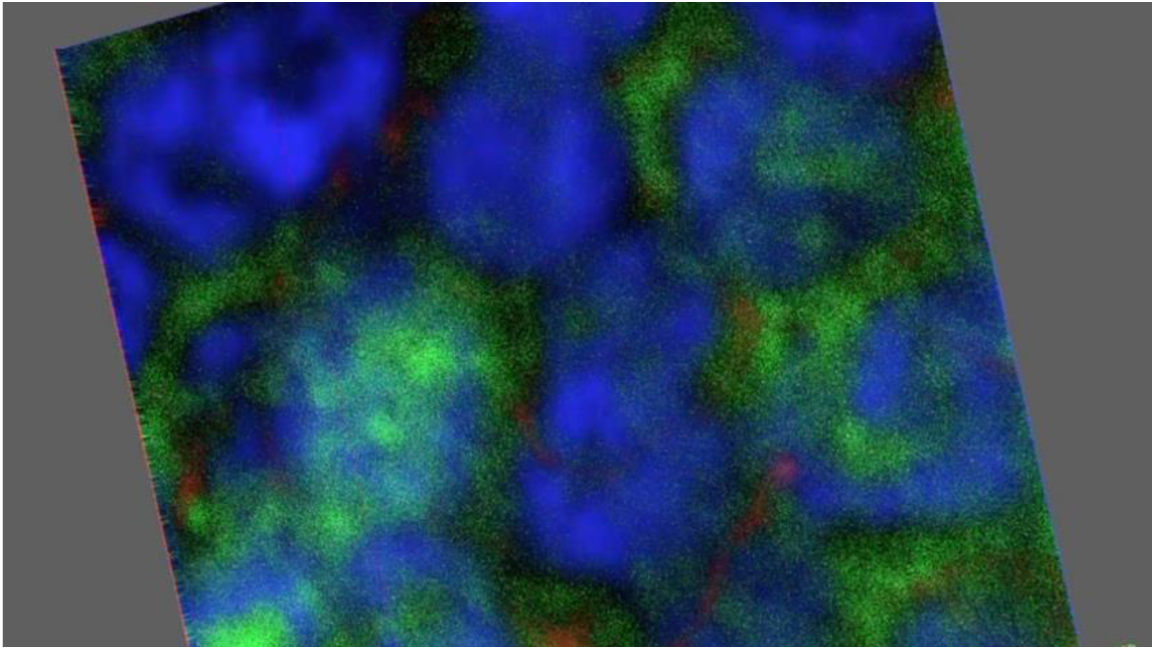
3D demonstration of Ago1a inside of *Buchnera* from a z-stack of confocal images. Arrows in the video point the *Buchnera* in focus. Blue represents DAPI staining of DNA; Red indicates phalloidin staining of F-actin for symbiosomal membrane around *Buchnera*; Green indicates Ago1a signal.

